# CovProfile: profiling the viral genome and gene expressions of SARS-COV-2

**DOI:** 10.1101/2020.04.05.026146

**Authors:** Yonghan Yu, Zhengtu Li, Yinhu Li, Le Yu, Wenlong Jia, Yiqi Jiang, Feng Ye, Shuai Cheng Li

**Author notes:** The authors contributed equally to the work (Yonghan Yu, Zhengtu Li, Yinhu Li, Le Yu). **Corresponding authors** Correspondence to Feng Ye and Shuai Cheng Li.

## Abstract

The SARS-CoV-2 virus has infected more than one million people worldwide to date. Knowing its genome and gene expressions is essential to understand the virus’ mechanism. Here, we propose a computational tool CovProfile to detect the viral genomic variations as well as viral gene expressions from the sequences obtained from Nanopore devices. We applied CovProfile to 11 samples, each from a terminally ill patient, and discovered that all the patients are infected by multiple viral strains, which might affect the reliability of phylogenetic analysis. Moreover, the expression of viral genes ORF1ab gene, S gene, M gene, and N gene are high among most of the samples. While performing the tests, we noticed a consistent abundance of transcript segments of MUC5B, presumably from the host, across all the samples.

## Introduction

Severe acute respiratory syndrome coronavirus 2 (SARS-CoV-2) has infected more than 1.2 million persons, and has led to more than 68,000 deaths, at the time of preparation of this manuscript [1–4]. The virus is an enveloped and single-stranded RNA betacoronavirus of ∼30k base-pairs (bps), which belongs to the family Coronaviridae [5, 6]. Since the year 2000, we have witnessed and experienced three highly widespread pathogenic coronaviruses in human populations — the other two are SARS-COV in 2002-2003, and MERS-CoV in 2012 [7]. All three viruses can lead to Acute Respiratory Distress Syndrome (ARDS) in the human host, which may cause pulmonary fibrosis, and lead to permanent lung function reduction or death [8–10]. The mechanisms of the severe cases due to SARS-CoV-2 remain unclear.

No drug is currently confirmed to be effective against SARS-CoV-2. Some vaccine candidates are under intensive studies [11]. A nucleotide analog RNA-dependent RNA Polymerase (RdRP) inhibitor called remdesivir has demonstrated effects on SARS-CoV-2 [12]. Hoffmann *et al.* suggested that TMPRSS2 inhibitor approved for clinical use can be adopted as a treatment option [13]. *In silico* methods have as well produced lists of potential drugs [14] and epitopes [15] worthy of further exploration. However, short of details on how the viral molecule compromises the host’s cells, it is hard to prioritize our exploration of the huge array of potential drugs and therapies. Analysis of the mechanism can provide us insights on the viral transmission as well as reveal therapeutic targets.

However, there are many pitfalls in sequencing the SARS-CoV-2 virus. Since the samples typically carry very low viral load, they are necessarily amplified through multiplex reverse transcription polymerase chain reaction (RT-PCR) before sequencing. Nanopore sequencing is typically used, in order to generate long reads that allow the detection of large genomic structural changes, viral recombination, and viral integration into the host genome [16]. They also support alternative splicing analyses of microorganisms. The approach has been used to sequence the dengue [17] and Zika viruses [18], as well as the SARS-CoV-2 [19].

This approach, however, has several problems. First, the long reads from Nanopore technology suffers from high error rates. Second, the sub-sequences can be amplified unevenly. Third, reads in the gene region amplified from the viral genome are indistinguishable from the gene transcripts. Finally, chimeric reads are frequent. These inaccuracies severely impact the study of viral expressions. Furthermore, while not inherent in the technology, we noticed that the problem could be further complicated by the existence of multiple viral strains.

Here, we developed a computational tool, CovProfile, to detect viral genomic variations, and profile viral gene expressions through data from multiplex RT-PCR amplicon sequencing on Nanopore MinION. To detect the genomic variations, we mapped the reads to SARS-CoV-2 reference genome (MN908947.3). To profile the viral gene expressions, we adopt a regularization method to find out the primer efficiencies. The tool was tested on the data obtained from the sputum samples in eleven severe COVID-19 patients. No structural variation was found, but all eleven patients appear to be infected by two strains. We found that the expression of viral genes ORF1ab gene, S gene, M gene, and N gene are high among most of the samples. While performing the tests, we noticed that MUC5B transcript segments to be consistently more abundant than segments of other genes in the host. The function of this gene may be associated with the severe symptom of abundant mucus in COVID-19 patients.

The discovery of multiple strains in all our samples has severe implications on the study of phylogeny, which rely heavily on SNPs (Single Nucleotide Polymorphisms). We identify at least two problems. First, coexistence of multiple strains might lead to inference of the wrong allele combinations, resulting in incorrect phylogeny and interpretation. Second, SNV calling might, in fact, be incorrect when mutations are located near the primer region. This problems are severe and requires more careful analysis, since phylogeny is essential in studying the spreading of the pandemic and evolution of the virus [20–23].

## Method and Materials

### RNA reverse transcription and Nanopore sequencing

Sputum samples were collected from the 11 patients and inactivated under 56°C with 30 minutes in accordance with WHO and Chinese guidelines [24, 25]. Then, the total RNA was extracted from the samples according to the protocol of RNA isolation kit (RNAqueous® Total RNA isolation Kit, Invitrogen, China), and RNA concentration was determined by Qubit (ThermoFisher Scientific, China). Based on 2 pools of primers (98 pairs of primers in total) (Supplementary table1), the entire genomic sequence of SARS-CoV-2 was amplified segmentally by reverse transcription. Then, the library was built by Nanopore library construction kit (EXP-FLP002-XL, Flow Cell Priming Kit XL, YILIMART, China), while the adapter and barcode sequences were also added to the samples. The samples were sequenced on the Oxford Nanopore MinIon.

### Data conversion and filtration

Fast5 format data was generated by the sequencer, and converted into fastq format with guppy basecaller (version 3.0.3). By applying NanoFilt (version 1.7.0) [26], we performed data filtration on the raw fastq data with the following criteria: read lengths should be longer than 100 bp after the removal of the adapter sequences, with an overall quality higher than 10. The clean data was kept for subsequent analysis.

### Data composition

A total of 1,804,043 clean reads were generated, with an average of 164,004.00±90,848.28 (Mean±SD) reads per sample (Fig 1). Aligning the reads to SARS-CoV-2 and human genomes respectively, the alignment ratio ranged from 20.14% to 99.74% on SARS-CoV-2 genome, and ranged from 0.11% to 72.12% on the human genome. Furthermore, each sample contained 71.24±35.36M data on average, while 61.29±42.13M and 8.87±9.34M data were aligned to SARS-CoV-2 and host genomes, respectively.

**Figure 1.**
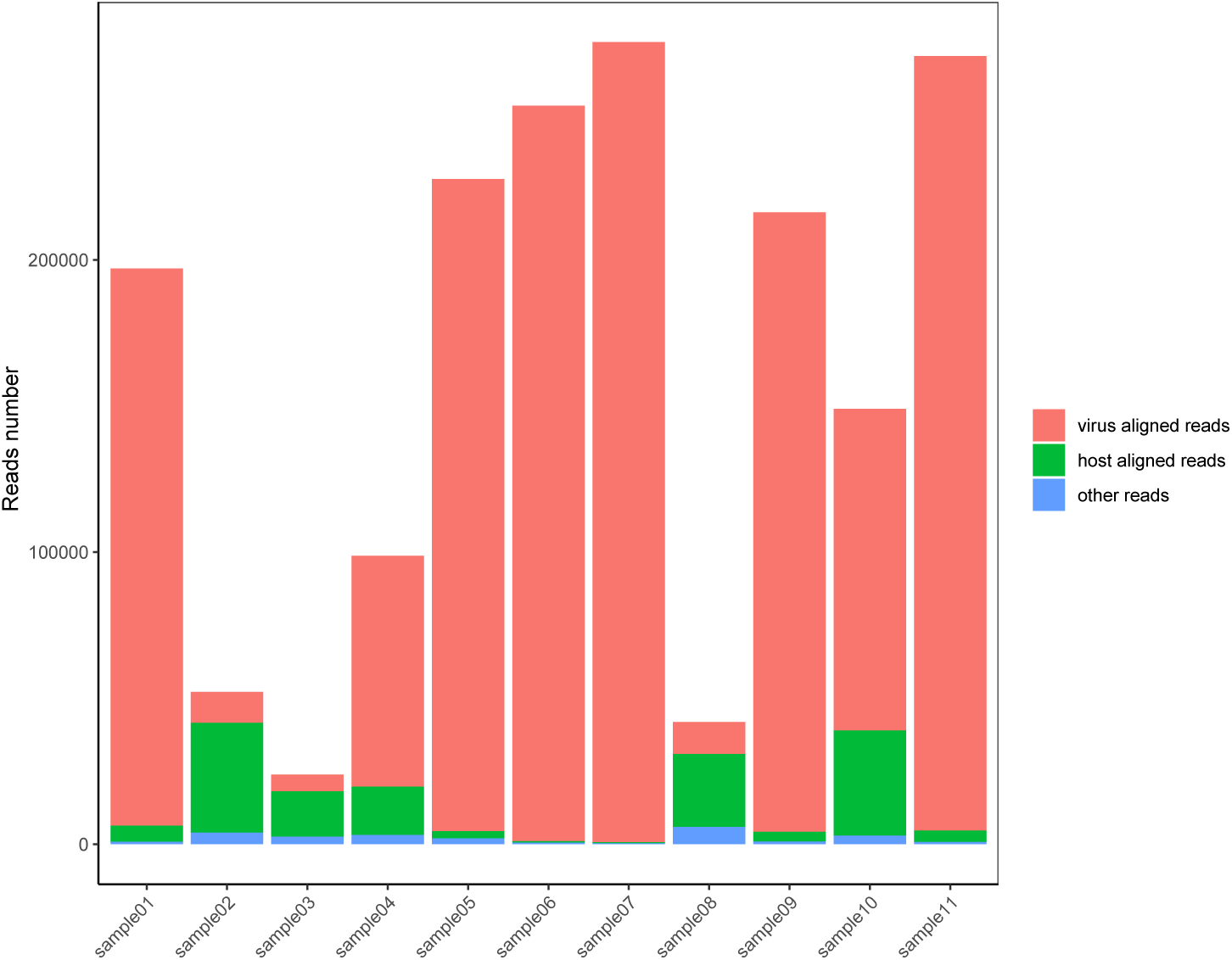
Compositions of the data sets. The heights of the columns represent the data composition. The data from the host, SARS-CoV-2 and other genomes are represented respectively with the colors pink, green and blue.

### Remove false-positive chimeric reads

Chimeric Nanopore reads formed by random connection of multiplex RT-PCR amplicons were split into multiple segments if the locations of their bilateral ends match corresponding locations of PCR primers. These chimeric reads were able to be split into several segments with their bilateral ends exactly matching the sites where the multiplex PCR primers are located, suggesting that they should be processed accurately to avoid false positive identification of virus recombinations or host integrations. We position the primers on both viral and host genome, and split the identified chimeric reads into segments corresponding to PCR amplicons. This method allowed us to salvage large amount of sequencing data, leading to more accurate alignment and higher coverage. Realignment were performed on the separated segments before analysis.

### Mutation detection

We aligned the clean data to SARS-CoV-2 genome (NCBI MN908947.3) with minimap2 [26] with default parameters for Oxford Nanopore reads. The aligned PCR amplicons were separated according to corresponding primer pool. With the separated alignment results, the genomic variations with average quality larger than 10 were called with bcftools (version 1.8). Mutations with less than 10 supported reads were filtered. To reduce the effects brought about by the PCR amplification, a variation is filtered if it was located within 10 bp upstream or downstream of the primer region within corresponding primer pool. The filtered mutations for different primer pools were then merged as the final mutations. Based on the gene information in SARS-CoV-2 reference genome, the final mutations were annotated by self-programming software.

### Estimating viral load and primer efficiency

Assume that there are *n* pairs of primer (*s*_*i*_, *t*_*i*_), with primer efficiency *e*_*i*_, 1 ≤ *i* ≤ *n*. Let the viral gene boundaries be denoted (*l*_*g*_, *r*_*g*_), where *g* indexes the genes. Denote the observed sequence depth captured by the primer pair (*s*_*i*_, *t*_*i*_) as *d*_*i*_.

We want to estimate the viral gene expression. Let the number of viral copies be denoted *v*, the number of transcripts of gene *g* be denoted *t*_*g*_, and the sequencing rate of amplified segment be denoted *α*. Then, we can have the following two approximations for *d*_*i*_.

If the primer *i* is fully covered by gene *g* (Fig1. *A*), then as the number of amplicons increases exponentially with the number of replication cycles we have

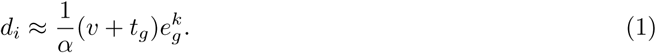

where *k* is the number of replication cycles.

Similarly, if primer pair *i* spans across gene boundaries; that is, it belongs to none of the gene transcripts (Fig1. B), then we have

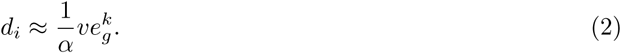

If some reads share two ends, where one end is a primer and the other end is at some gene boundary (Fig 1D, Fig 1E), it may indicate that these amplicons are the result of linear amplification. Denote the number of reads *r*_*i,g*_ with two endpoints as (*t*_*i*_, *e*_*g*_), and the number of reads *r*_*g,i*_ with two endpoints as (*l*_*i*_, *s*_*g*_), then

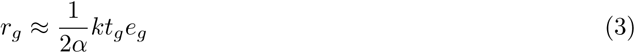

If a gene *g* is within some primer target region (Fig2. G), then the sequenced fragment can be expected to exactly cover the gene. Let the number of observed reads span exactly the whole gene *g* as *r*_*g*_, then

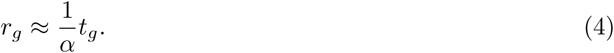

**Figure 2.**
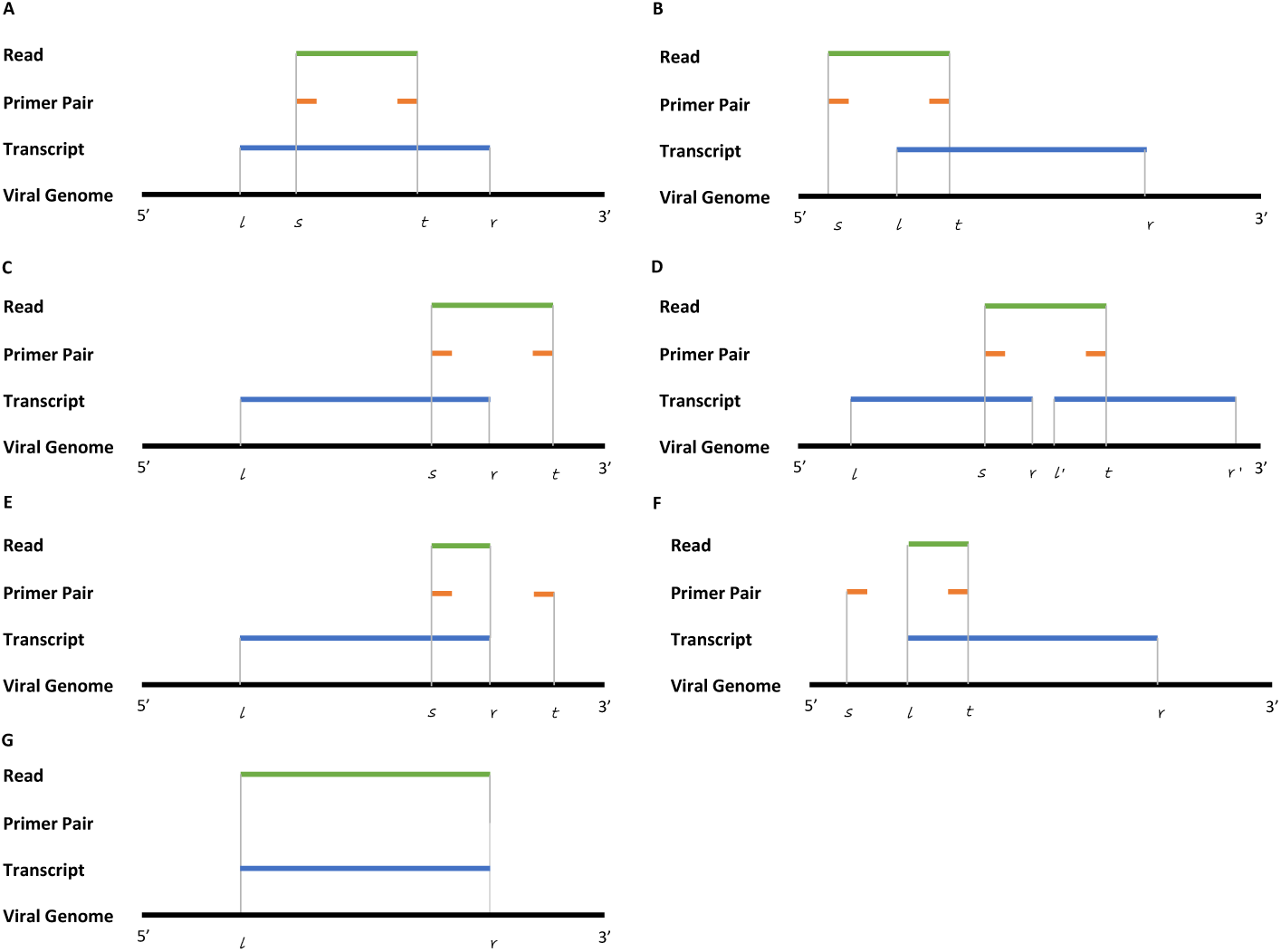
Relations between viral gene, primer pair and sequence. **A**: Primer pair spans fully on gene with sequence aligned with primers. **B**: Primer pair spans across gene boundaries at 5’ with sequence aligned with primers. **C**: Primer pair spans across gene boundaries at 3’ with sequence aligned with primers. **D**: Primer pair spans across two gene with sequence aligned with primers. **E**: Primer pair spans across gene boundaries at 5’ with sequence aligned with gene boundaries at 3’ and primer at 5’. **F**: Primer pair spans across gene boundaries at 3’ with sequence aligned with gene boundaries at 5’ and primer at 3’. **G**: Sequence aligned with gene boundaries at both 5’ and 3’.

To find the expressions of the viral genes, we want to minimize the following objective function based on the entities defined above

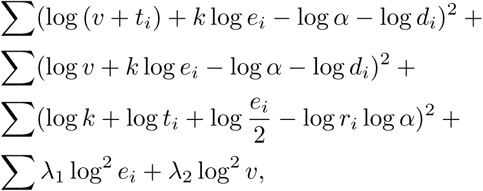

where λ_1_ and λ_2_ are two regularization parameters, and *k* is a weight factor (we set *k* to 21 according to our experiments). The viral load *v*, primer efficiencies *e*_*i*_, and expressions *t*_*g*_ which minimize the objective equation are our estimation for these parameters.

### Calculate viral gene expression

To calculate the viral gene expression, the duplication generated during PCR were removed based on the estimated primer efficiencies. The gene expression can be calculated as the sum of the base depth of each gene divided by its length.

### Capture gene product from host

To capture the gene products of the host, reads and primers were aligned to the hg38 transcript database derived from UCSC refGene annotation. Primer alignment were performed with BLAT [27]. Regions where the depth of reads alignment are equal or larger to 5 are screened. Reads with endpoints matching the screened primer alignment were treated as PCR duplications. Due to mismatches in primer to transcript alignment, primer efficiencies may change when capturing gene product. Thus, the recognized PCR duplication were removed and one copy of those reads were kept. The gene expression from host can then be estimated as the sum of the base depth of each transcript divided by its length.

## Results

### Distribution of SNPs in SARS-CoV-2 genome

After aligning the sequencing data to the SARS-CoV-2 genome, we detected the mutations of SARS-CoV-2 in the 11 samples (Figure 3). Based on these mutations, we discovered a total of 48 SNPs (Supplementary table2); 44 of them located on the genetic regions, including gene ORF1ab, S, N, M, ORF6 and ORF8. Gene ORF1ab contained 28 SNPs in 11 samples, and 20 of them were non-synonymous mutations. Non-synonymous mutations also occupied a large part of SNPs in the gene S, N, M and ORF8: 7 of 10 SNPs in gene S, 1 of 3 SNPs in gene N, 1 of 1 SNPs in gene M, and 1 of 1 SNPs in gene ORF8 were non-synonymous mutations. Since most of the SNPs were located in only 1 sample, we deduced that the functions of the discovered SNPs need to be further verified according to lab experiments. Surprisingly, we also found two SNPs (loci 8,782 and 28,144), which were important SNP loci identified in recent phlogenetic analysis [20], that frequently demonstrates heterogeneous signals, suggesting the existence of at least two different virus strains in each host. However, we did not find creditable InDels (Insertions and Deletions), or structural variations, viral-host integration during the mutation analysis.

**Figure 3.**
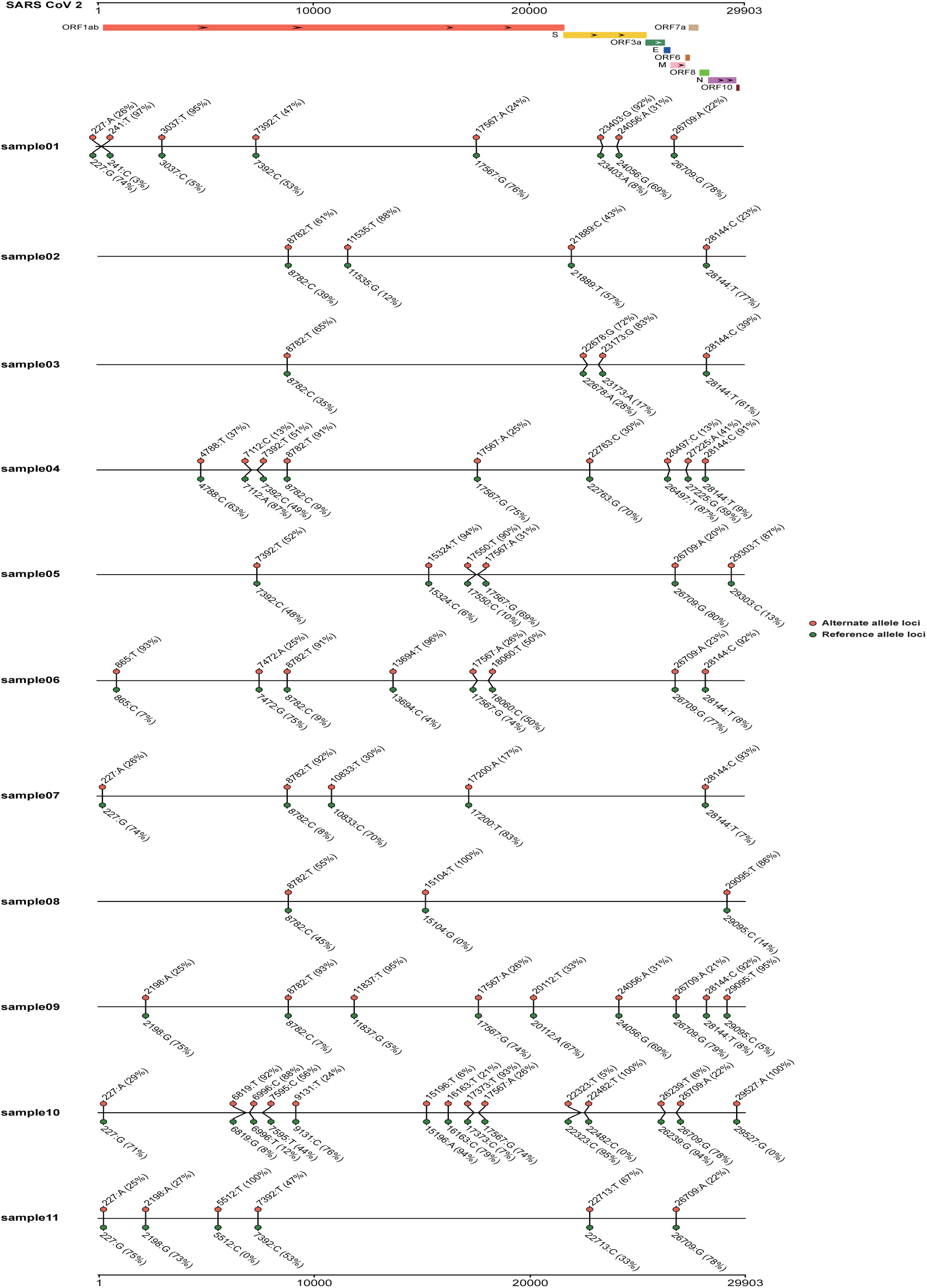
Distributions of SNPs among the 11 COVID-19 samples. The locations of genes in SARS-CoV-2 are displayed on top of the plot, with the distributions of SNPs for the 11 samples. The genotype of SNP and reference were list above and below the line respectively, with the percentages of their supported reads.

### Evaluations of the amplification efficiencies for the primers

Since 98 pairs of primers from two pools were designed for the enrichment of SARS-CoV-2 genome before sequencing, we discovered that different parts of SARS-CoV-2 genome exhibited different coverage depths. We then calculated the amplification efficiencies for the primers with our proposed method. We discovered that the primers located at the 5’ and 3’ ends of the SARS-CoV-2 genome exhibited higher amplification efficiency (>1.2, Figure 4), including primer01 and primer84-primer93. With the genomic locations of the primers, we know that the genomic sequences within the ranges 24-410 and 25,280-28,464 can be amplified with more sequencing data. On the other hand, the remaining primers exhibited slightly lower efficiencies, within the range of 1.0-1.2. For primers NO. 64, 70, 83 and 85, their amplification efficiencies were as low as 1, which indicates that the target regions of these primers were not amplified. Based on the primer efficiencies, the effect of data bias which was induced by the PCR experiments can be attenuated from the original data, and they can be further applied for the normalization of viral gene expression under different primers.

**Figure 4.**
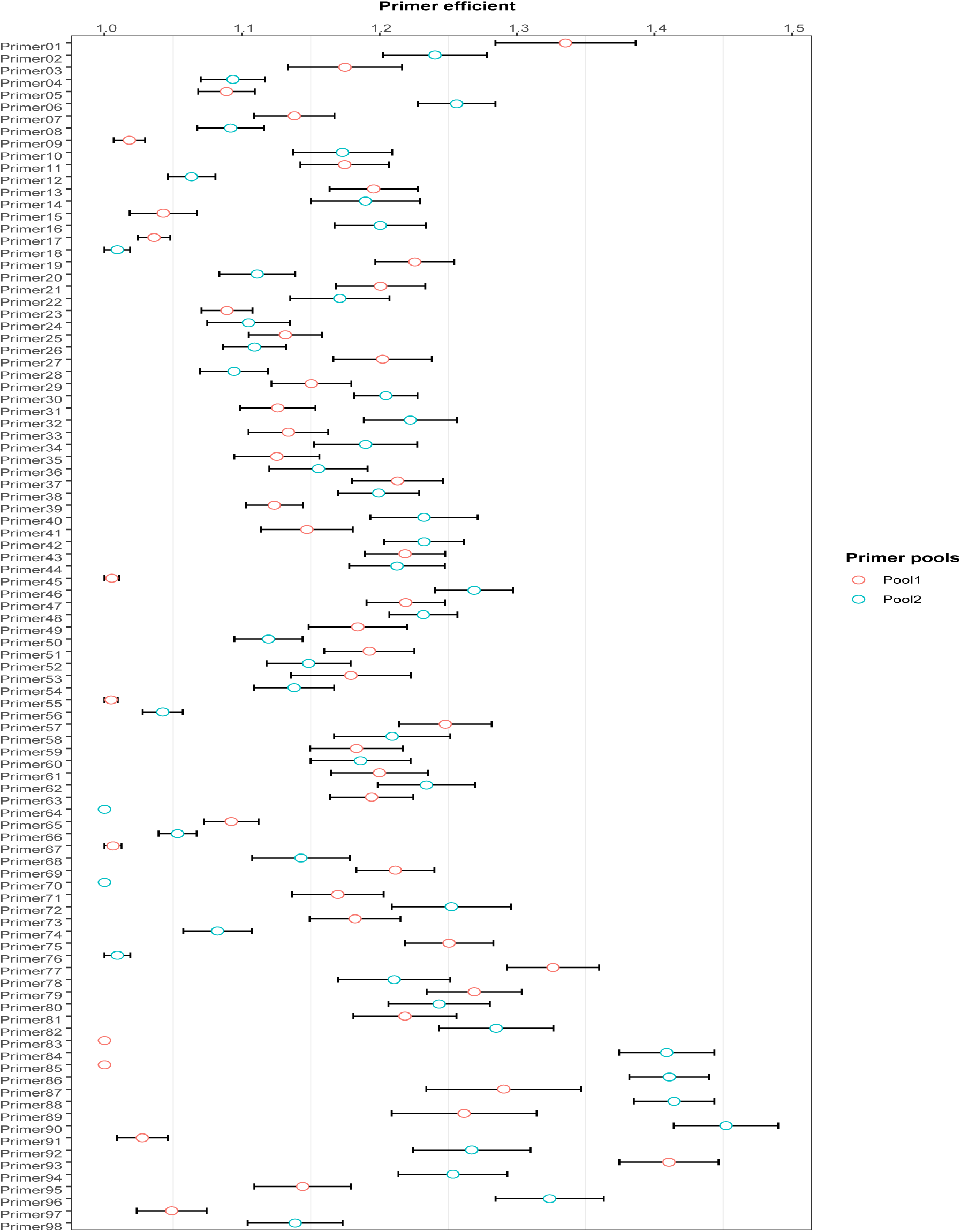
Amplification efficiencies for the 98 pairs of primers. The amplification efficiencies calculated for the primers, and their distributions in 11 samples. The primers in pool1 and pool2 are represented using red and blue markers respectively.

### Normalized coverage depth on SARS-CoV-2 genome

Due to the amplification discrepancy among the primers, the sequencing data was unevenly distributed along the SARS-CoV-2 genome. After aligning the sequencing data to the SARS-CoV-2 genome, we found that the coverage depths of some genomic regions reached over 8000x in some samples, but some regions have coverage depths less than 10x (Figure 5). In order to minimize the data imbalance induced by PCR amplification, we performed data normalization based on the primer efficiency and their locations on SARS-CoV-2 genome, so that the normalized data could reflect the actual virus reproduction in the host. Based on the algorithm illustrated in the method, we obtained the normalized sequencing depth of 11 sample data.

**Figure 5.**
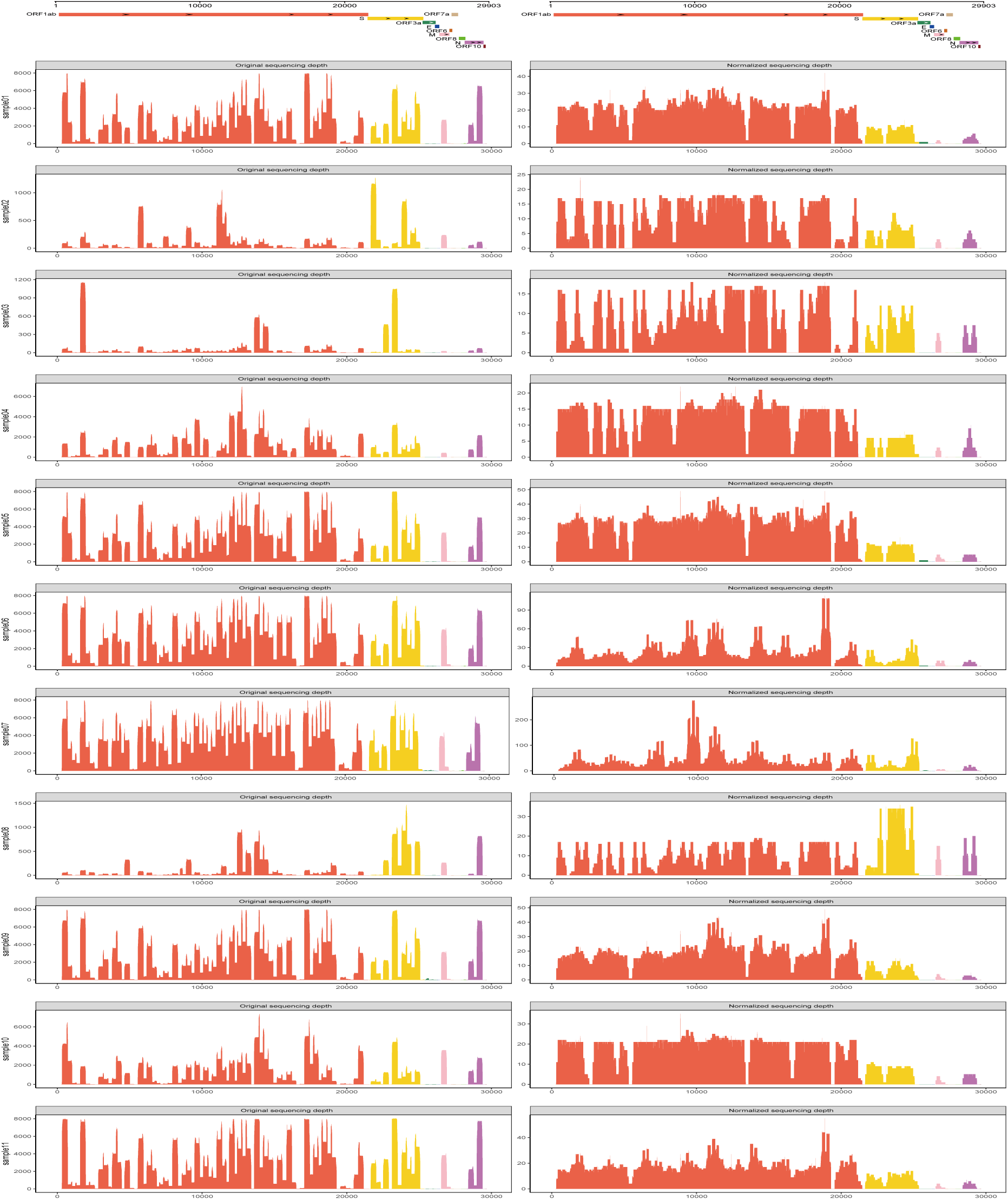
Original and normalized coverage depth for the 11 samples. The locations of the genes in SARS-CoV-2 are displayed on top of the plot, and marked with different colors. Each sample contains two plots, respectively representing the original (left) and normalized coverage (right) depths for the bases along the SARS-CoV-2 genome. The coverage depth are colored by the genes in SARS-CoV-2.

**Figure 6.**
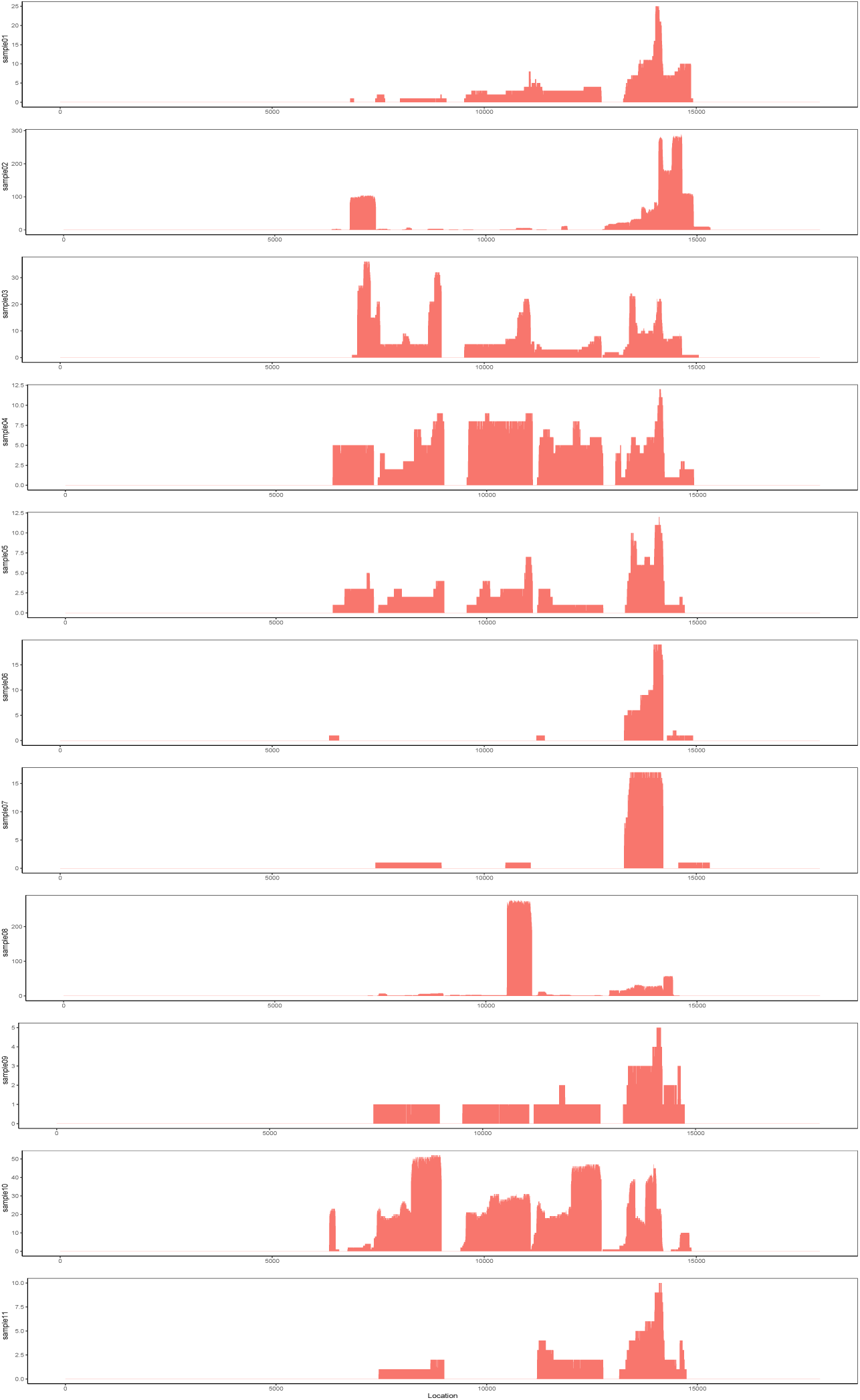
Normalized coverage depth on MUC5B gene in the 11 samples. In each plot, the X and Y coordinates respectively represent the length of MUC5B and the normalized coverage depth along the MUC5B gene.

After data normalization, the overall coverage depth along the SARS-CoV-2 genome was relatively even, and less than 300x. In addition, we discovered that the ORF1ab gene, S gene, M gene, and N gene exhibited higher coverage depth in the samples, which might indicate higher expression (Figure 5). These genes are related to host cell recognition, amplification and virus particle assembly, so we speculate that the results might explain the severe SARS-CoV-2 virus infection in the patients. Since gene E, ORF6, ORF7a, ORF7b, ORF8 and ORF10 are less than 400bp in length, their transcripts might not have been captured by the primers, and hence their expressions cannot be estimated.

### Highly expressed genes from the host

We noticed the existence of human genome in the samples from an earlier study. We similarly analyze the corresponding genes and their expression levels. Aligning these data to human transcript database, we identified the segments with possibly human gene transcripts. The coverage depths of these genes were normalized based on the primer efficiencies (Supplementary table3). Only several genes are listed within the top 5% in more than five samples according to the normalized depth. They include the genes MUC5B, TCAF12, KRT12 and *etc*. MUC5B is the only gene consistently within the top 5% across all the samples. More than 50% of the MUC5B gene region is covered (Figure 5). However, the region with high coverage in MUC5B gene is inconsistent among the samples, perhaps due to data scarcity.

## Conclusion and Discussion

In this study, we implemented a software tool, CovProfile, to analyze multiplex RT-PCR on Nanopore sequencing data, and studied the sputum samples of COVID-19 patients. Our results show enrichment of PCR amplicons which in total cover 99.7% of SARS-CoV-2 virus genome, confirming the reliability of the multiplex RT-PCR method in identifying COVID-19 infection.

The data contained large amount of chimeric reads formed by unintended random connection of multiplex PCR amplicons, making up 1.69% of total sequencing reads. Similar observation has been reported that these chimeric reads can be formed during library preparation and sequencing, and lead to the cross barcode assignment errors in multiplex samples [28, 29]. Our analysis of the host genome expression reveals several genes that are consistently activated in the majority of samples after eliminating the PCR effects. The most abundant gene is the MUC5B, which encodes a member of the mucin protein family. The family of proteins makes up the major component of mucus secretions in normal lung mucus [30]. MUC5B has been reported to be highly expressed in lung disease, such as chronic obstructive pulmonary disease (COPD). A recent research demonstrated potential abundance of gene MUC1 and gene MUC5AC in the sputum of COVID-19 infected patients [31]. Its up-regulated transcription could cause mucociliary dysfunction and enhances lung fibrosis [32–34], thus might be relevant to the recent report of large quantity of mucus found in the lung of COVID-19 infected patients. Another activated host gene is the TCAF2, which encodes transient receptor potential cation channel-associated factor. It could bind to the TRPM8 channel and promote its trafficking to the cell surface. Previous study reported that TCAF2 significantly increases the migration speed of prostate cancer cells when it co-expresses with TRPM8 [35]. Other enriched genes might also break the cellular homeostasis, including the nucleoredoxin-like gene NXNL2, tyrosine kinases FER, and so on. Further studies of these genes could help researchers gain insights into the dysregulation of host cells after the SARS-CoV-2 infection. We note that these genes are extracted based on a very limited amount of data, and the conclusions can be different with a large data set.

The viral expressions we uncovered show recurrent heterogeneous SNVs, not necessarily from the same loci, on every one of our 11 samples. This has severe implications on phylogenetic analysis on SARS-CoV-2, as the heterogeneous loci might be mis-linked during assembly.

This observation led us to believe that the coexistence of multiple virus strains may be a common phenomena. To verify this assumption, we downloaded all 4,075 SARS-CoV-2 strains available from GISAID at the time of preparation of this manuscript to analyze the complexity of co-occurrences of SNVs within the dataset. We consider the occurrence of the genotypes on two different loci. If every possible combination of the genotypes on the loci pair exists in at least one strain in the dataset, we say that the loci is *complex*. A complex loci is difficult to explain in phylogeny, but may occur by chance. To remove this possibility, instead we perform an analysis based on *SNP cliques*. We model a single SNP locus as a vertex and create an edge between a loci pair if and only if each of the four combinations of the genotypes of the pair is detected in at least one strain within the dataset. In this case, the existence of a large clique will be intractable to explain using phylogeny, as the chances become diminishingly small with the size of the clique.

From the dataset, we were able to uncover a partial clique of 24 vertices and 138 edges. Two maximal 8-cliques and 14 maximal 7-cliques from the partial cliques are shown in Fig 7. These cliques suggest fast mutations on their loci. On the other hand, genomic recombination between the strains is not detected with the software package RDP4 [36]. Hence, we believe the cliques would be best explained by the coexistence of multiple SARS-CoV-2 strains in a host. In this case, different loci from different viral strains might have been mixed and reported as a single SARS-CoV-2 genome, thus resulting in the prevalence of complex loci. To verify this hypothesis, we examined SARS-CoV-2 Nanopore sequencing data under accession SRR11347377, SRR11313280, SRR13313419, SRR13313494, SRR11140745, SRR11140747, SRR11140749 and SRR11140751, which were collected from San Diego, Wuhan and Wisconsin Madison, respectively. Then, credible evidence of multiple viral strains were identified (See Supplementary Table 4). We also discovered that loci 8782, 26144 and 28144, which were important loci in recent SARS-CoV-2 phylogenetic analysis [20], were involved in some cliques of over 5 loci (Supplementary Figure 1).

**Figure 7.**
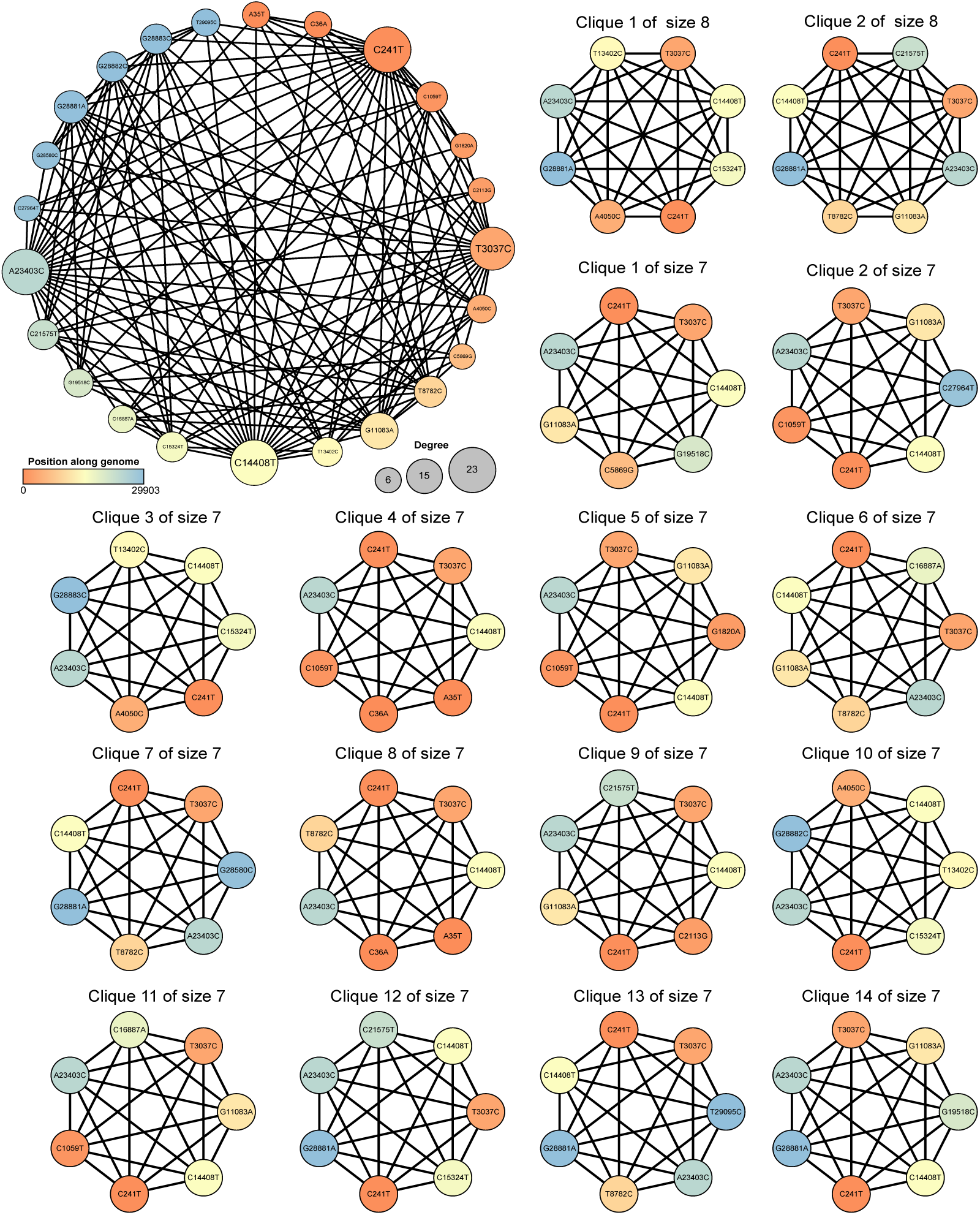
Maximal SNP *k*-clique with *k* > 6.

In closing, we expect CovProfile to continue to help in detecting genomic variation on transcriptional profile of SARS-CoV-2 infection from multiplex Nanopore sequencing data. The tool decomposes and realign the chimeric reads that commonly exist in the multiplex Nanopore sequencing, and this will greatly promote data usage. The method also provides robust estimation of primer efficiency through a multi-parameter model. We hope the viral and host gene expressions we reported could help in the development of diagnosis and therapy strategies of SARS-CoV-2 infection candidates.

## Data Availability

The raw sequence data reported in this paper have been deposited in the Genome Sequence Archive [37] in BIG Data Center [38], Beijing Institute of Genomics (BIG), Chinese Academy of Sciences, under BioProject PRJCA002503 with accession ID CRA002522 that are publicly accessible at https://bigd.big.ac.cn/gsa. All other data are available from the authors upon reasonable request.

## Code Availability

CovProfile used in the main analysis pipeline are available at https://gitlab.deepomics.org/yyh/covprofile. All other code are available from the authors upon reasonable request.

## Acknowledgments

We thank every participants contributing to this work.

## Author Information

These authors contributed equally: Yonghan Yu, Zhengtu Li, Yinghu Li and Le Yu.

### Contributions

SCL conceived the idea.

SCL and FY supervised the project.

SCL and YY designed the algorithms

SCL, WJ, YY and YL discussed and designed the experiments.

YY and YL implemented the code and conducted the analysis.

YJ conducted the clique analysis.

ZL and LY prepared the data.

SCL YY WJ and YL drafted the manuscript.

SCL, FY revised the manuscript.

All authors read and approved the final manuscript.

## Ethics declarations

### Ethics oversight

This study was approved by the First Affiliated Hospital of Guangzhou Medical University, and the sample and data collection procedures were conducted in accordance with the principles expressed in the Declaration of Helsinki. All patients provided written informed consent, and volunteered to receive investigation for scientific research.

