## Supplementary figures and images for "CovProfile: profiling the viral genome and gene expressions of SARS-COV-2"

### Supplemental Fig 1

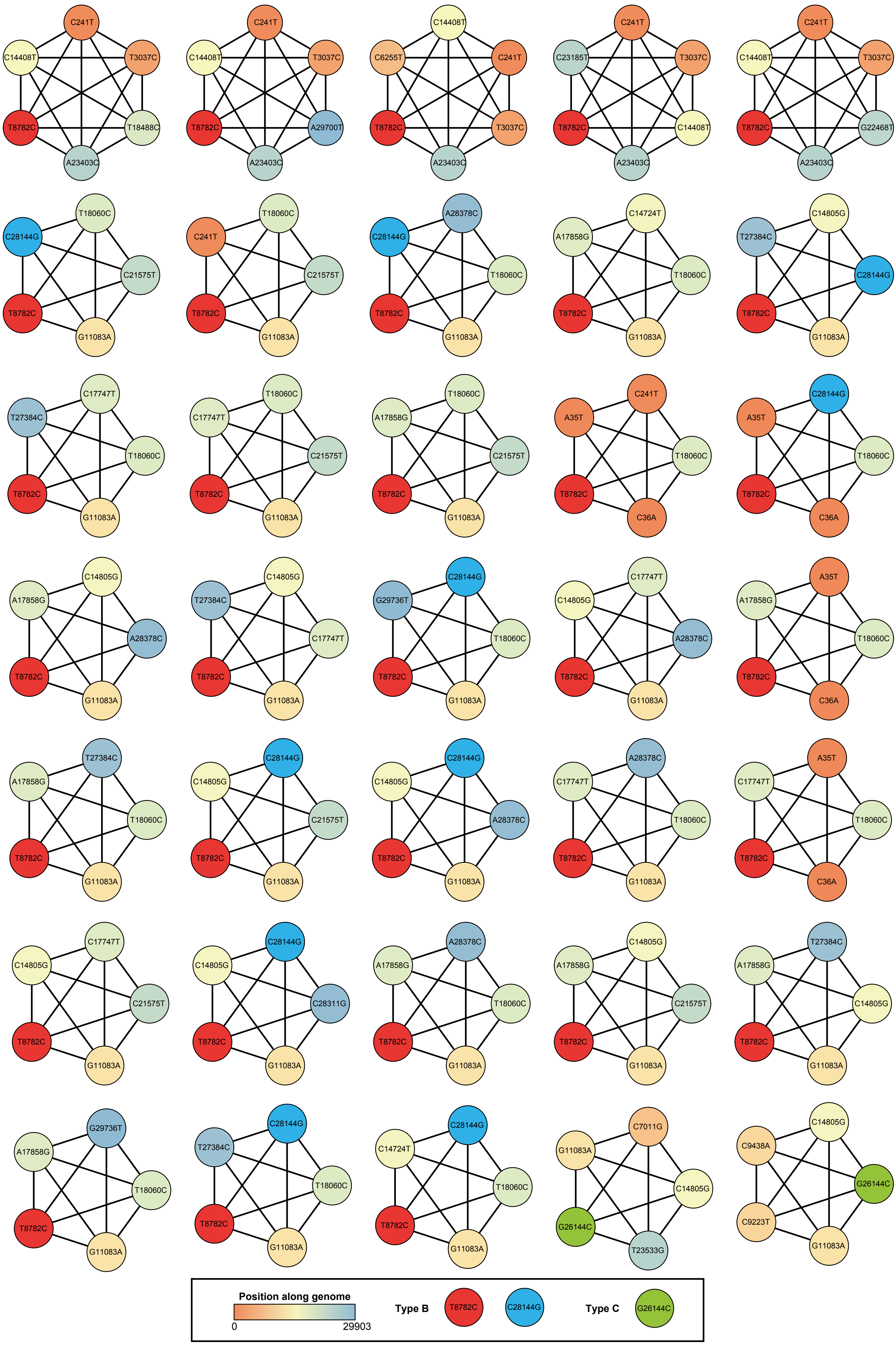
